# Chord: Identifying Doublets in Single-Cell RNA Sequencing Data by an Ensemble Machine Learning Algorithm

**DOI:** 10.1101/2021.05.07.442884

**Authors:** Ke-Xu Xiong, Han-Lin Zhou, Jian-Hua Yin, Karsten Kristiansen, Huan-Ming Yang, Gui-Bo Li

## Abstract

High-throughput single-cell RNA sequencing (scRNA-seq) is a popular method, but it is accompanied by doublet rate problems that disturb the downstream analysis. Several computational approaches have been developed to detect doublets. However, most of these methods have good performance in some datasets but lack stability in others; thus, it is difficult to regard a single method as the gold standard for each scenario, and it is a difficult and time-consuming task for researcher to choose the most appropriate software. To address these issues, we propose Chord which implements a machine learning algorithm that integrates multiple doublet detection methods. Chord had a higher accuracy and stability than the individual approaches on different datasets containing real and synthetic data. Moreover, Chord was designed with a modular architecture port, which has high flexibility and adaptability to the incorporation of any new tools. Chord is a general solution to the doublet detection problem.

## Introduction

Recently, the development of high-throughput single-cell RNA sequencing (scRNA-seq) has provided unprecedented convenience for dissecting the cellular heterogeneity of tissues(Wu and Zhang, 2020). In contrast to traditional transcriptomic studies using bulk RNA sequencing for only a virtual average gene expression of whole bulk samples, profiling transcriptomes at the single-cell resolution has enabled researchers to recognize the molecular characteristics of all cell types at one time and acquire a better insights into physiology, biological development and disease(Potter, 2018). Among the current state-of-the-art technologies of high-throughput scRNA-seq, droplet-based technologies are currently commonly employed as an unbiased solution of single-cell transcriptomics(Prakadan, Shalek and Weitz, 2017). However, these microfluidic methods often encounter the problem of doublets, where one droplet may contain two or more cells with the same barcode during the distribution step of isolating single cells. Then the droplet of doublets is counted as a single cell in the data forming technical artefacts (Wolock, Lopez and Klein, 2019). According to the composition of doublets, doublets can be divided into two major classes: homotypic doublets, which originate from the same cell type, and heterotypic doublets which arise from distinct transcriptional cells generating an artificial hybrid transcriptome(McGinnis, Murrow and Gartner, 2019; Wolock, Lopez and Klein, 2019). Compared to homotypic doublets, heterotypic doublets are considered to have more impact on downstream analyses including dimensionality reduction, cell clustering, differential expression and cell developmental trajectories(Bernstein, Fong, Lam, Roy, Hendrickson and Kelley, 2020; Xi and Li, 2020). To reduce the number of doublets in experiments, decreasing the concentration of loaded cells is an effective control measure to obtain a lower doublet rate, but this approach also reduces the number of captured cells and dramatically increases the cost per sample(Bernstein, Fong, Lam, Roy, Hendrickson and Kelley, 2020; Zheng, Terry, Belgrader, Ryvkin, Bent, Wilson, Ziraldo, Wheeler, McDermott, Zhu et al., 2017). Thus, instead of avoiding doublets, several existing experimental techniques can be applied to identify doublets, such as the cell hashing method using oligo-tagged antibodies as an orthogonal information(Stoeckius, Zheng, Houck-Loomis, Hao, Yeung, Mauck, Smibert and Satija, 2018), MULTI-seq using lipid-tagged indices(McGinnis, Patterson, Winkler, Conrad, Hein, Srivastava, Hu, Murrow, Weissman, Werb et al., 2019) and demuxlet using natural eetic variations(Kang, Subramaniam, Targ, Nguyen, Maliskova, McCarthy, Wan, Wong, Byrnes, Lanata et al., 2018). However, there are inherent limitations to these experimental techniques used for doublet detection. First, these methods require their own sepcial experimental operations and additional costs, so they are not suitable for existing scRNA-seq data. Second, these techniques are only experimentally label doublets from different samples but ignore the kind of doublet generated by cells from the same sample or individual. Therefore several computational approaches have been developed to detect doublets in common scRNA-seq data, including these already generated data(Xi and Li, 2020). However, their results vary greatly, and there are noticeable differences even in the top-performing methods which were demonstrated in a benchmarking study(Xi and Li, 2020), so there is still a larger challenge in terms of accuracy due to the low concordance between individual methods and suboptimal accuracy of each method. In addition, in view of the unique characteristics brought by these specific mathematical algorithms and applicability to different scenarios, no method is the gold standard for each scenario; thus, it is a difficult and time-consuming task for researchers to choose suitable software.

To address these unmet needs, we propose Chord which implements an ensemble algorithm that aggregates the results from multiple representative methods to accurately identify doublets. The ensemble algorithm is a widely used technique of machine learning(Dietterich, 2000) which can boost the accuracy of somatic mutation detection(Fang, Afshar, Chhibber, Mohiyuddin, Fan, Mu, Gibeling, Barr, Asadi, Gerstein et al., 2015), culprit lesion identification(Al’Aref, Singh, Choi, Xu, Maliakal, van Rosendael, Lee, Fatima, Andreini, Bax et al., 2020) and so on. Compared to the individual methods, Chord was demonstrated an improved doublet detection accuracy and stability across different datasets of real and synthetic data. Moreover, Chord was designed with modular architecture port that is highly flexible and adaptable to the incorporation of any number of new tools. The open source Chord package in R is available through GitHub (https://github.com/13308204545/Chord).

## Results

### 1. The Chord workflow for accurate and robust ensemble algorithm-based doublet detection

The computational approaches to detect doublets in scRNA-seq data are grouped into two categories. The first strategy of one category uses the distance between simulated artificial doublets and the observation cells to identify doublets, for example, DoubletFinder(McGinnis, Murrow and Gartner, 2019) adopts this strategy to handle the doublet detection task as a binary classification problem. The second strategy used by cxds in the scds(Bais and Kostka, 2020) package is based on co-expressed ‘marker’ genes that are not `simultaneously` expressed in the same singlet cell but can appear in doublet cells. However, the performance of existing computational doublet detection approaches varies greatly including overall detection accuracy, impacts on downstream analyses and computational efficiency(Xi and Li, 2020), which shows much room for improving the performance of the doublet detection method. Here, we describe a new strategy based on an ensemble algorithm of machine learning for doublet identification. Our approach, Chord, integrates four representative computational doublet detection methods, DoubletFinder(McGinnis, Murrow and Gartner, 2019), DoubletCells(Lun, McCarthy and Marioni, 2016), bcds and cxds(Bais and Kostka, 2020), in R environment (Table S1), to employ these enhancements to improve doublet detection (Figure 1A).

**Figure 1.**
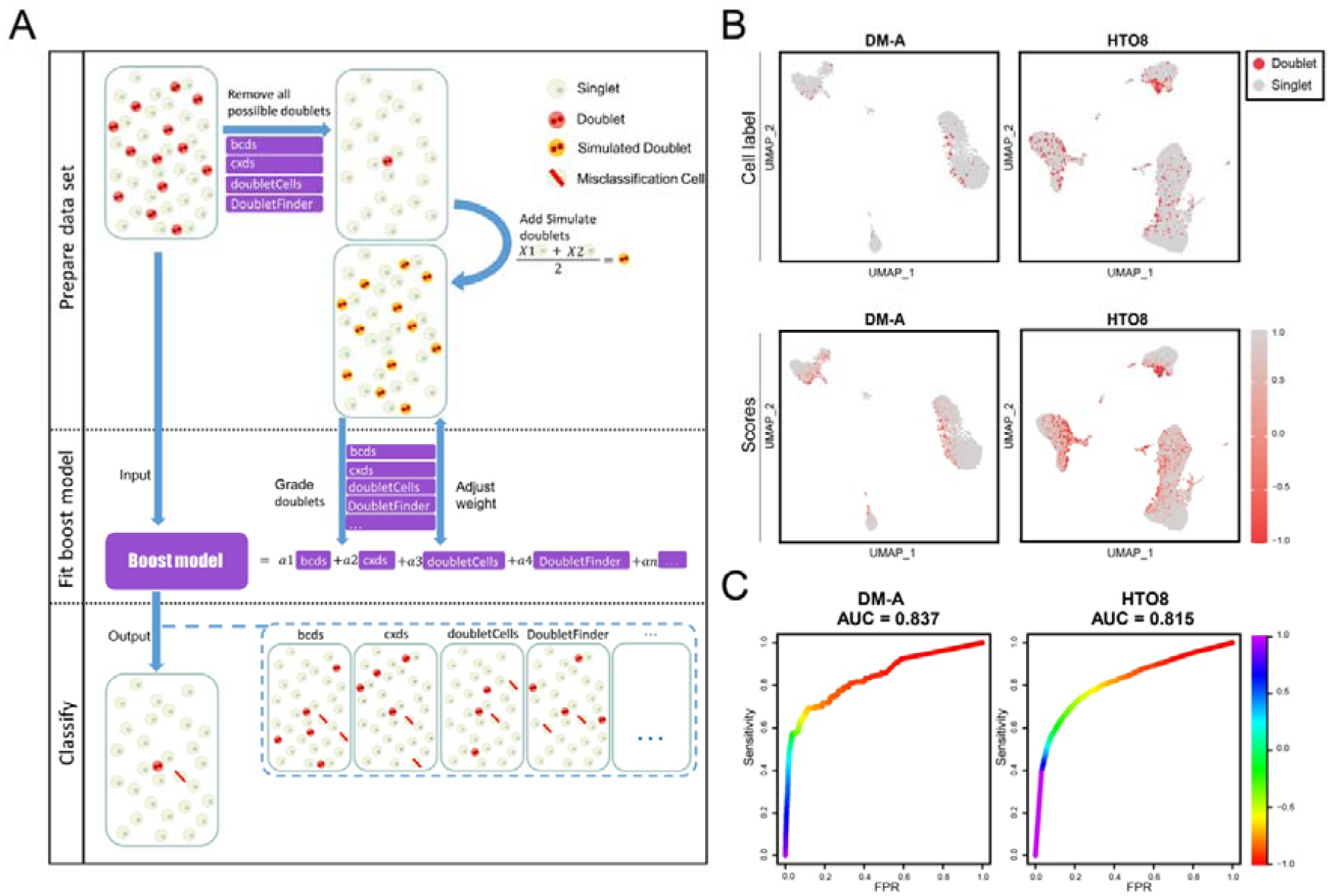
Chord overview and its performance on the test DM-A and HTO8 tests. (A) Schematic outline of the Chord workflow. `First`, preliminarily predicted doublets are filtered using bcds, cxds, doubletCells and DoubletFinder, `and` then the processed `dataset is randomly` sampled to generate simulation doublets `that` are added to `the` training `dataset`. The second step is to fit the weights of the integrated methods through the AdaBoost algorithm on `the` training `dataset`. In the third step, the ensemble model `is used` to evaluate the original expression matrix and the doublets are identified by the expectation threshold value. (B) UMAP was embedded for the DM-A and HTO8 `tests` with experimental `doublet` labels. The doublets are shown in red, and the singlets are shown in grey. The doublet prediction scores of Chord were visualized on the UMAP plots for the DM-A and HTO8 `tests`. The DM-A `dataset` was from `human peripheral blood mononuclear cell (PBMC)` samples using the experimental demuxlet method to annotate doublets(Kang, Subramaniam, Targ, Nguyen, Maliskova, McCarthy, Wan, Wong, Byrnes, Lanata et al., 2018). The HTO8 dataset was from the samples of cell lines using twelve barcoded antibodies to mark and label doublets(Stoeckius, Zheng, Houck-Loomis, Hao, Yeung, Mauck, Smibert and Satija, 2018). (C) The `ROC` curves of Chord were `drawn` for the DM-A and HTO8 `tests` using `the` R package PRROC(Grau, Grosse and Keilwagen, 2015). `The AUCs` for the DM-A and HTO8 datasets were 0.837 and 0.815`, respectively`.

The Chord workflow is composed of three main steps (Figure S1A). The program first roughly estimates the doublets of the input droplet data according to the four built-in methods to filter out the likely doublets from the original data before simulating artificial doublets. Therefore, we proposed a key step and an adjustable parameter called overkillrate to `preliminarily delete doublets`. Selecting this parameter could improve the accuracy of the program (STAR Methods; Figure S1A). Next, a simulation training set is generated from quality singlet data after removing these doublets. Finally, the AdaBoost algorithm was adopted to integrate these doublet detection methods to model training. Then the doublet scores output was calculated by the AdaBoost model for the input droplets data.

To determine whether the ensemble algorithm improves the performance of doublet detection, `we first`, evaluated these methods on ground-truth scRNA-seq datasets `that` label doublets using the experimental strategies demuxlet(Kang, Subramaniam, Targ, Nguyen, Maliskova, McCarthy, Wan, Wong, Byrnes, Lanata et al., 2018) and Cell Hashing(Stoeckius, Zheng, Houck-Loomis, Hao, Yeung, Mauck, Smibert and Satija, 2018). Ground-truth comparisons illustrate that Chord detects most doublets (Figure 1B, 1C). Using receiver operating characteristic (ROC) curve analysis and precision-recall(PR) curve analysis, the performance results for each dataset and `the` average across datasets are presented in Table 1 and supplementary Table S3. We find that our method Chord `performs` well on the HTO8 and DM-A datasets. Chord achieves the highest areas under the ROC curves (AUCs) in the identification results of the HTO8 and DM-A datasets (0.815 and 0.837, respectively), and Chord achieves the second highest area under the PR curve (AUPRC) value on the `HTO8 dataset` (0.530)`, which was` only lower than that of cxds (0.543), while Chord had `a` higher AUPRC value (0.345) than cxds (0.273) on the DM-A dataset (Figure S1B).

**Table 1.**
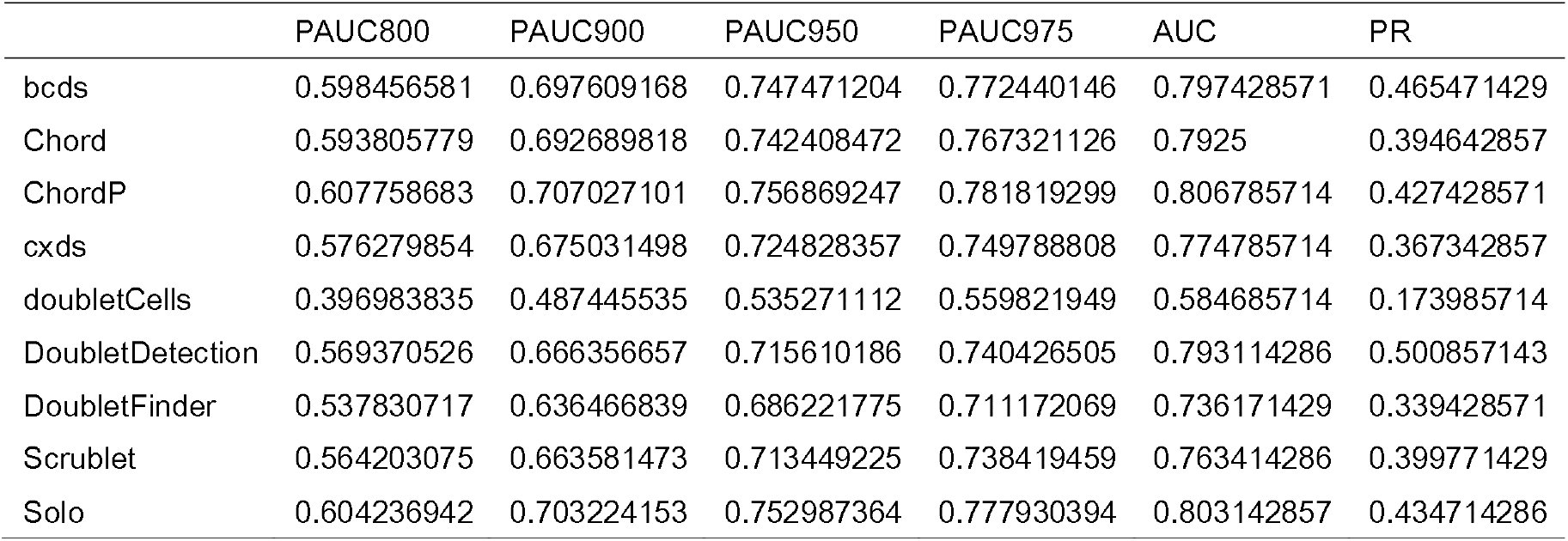
Comprehensive performance of each method in real-world scRNA-seq datasets with experimentally annotated doublets. The average performance of various methods in all the test datasets. The indicators are the pAUC800, pAUC900, pAUC950, pAUC975, AUC and AUPRC.

|  | PAUC800 | PAUC900 | PAUC950 | PAUC975 | AUC | PR |
| --- | --- | --- | --- | --- | --- | --- |
| bcds | 0.598456581 | 0.697609168 | 0.747471204 | 0.772440146 | 0.797428571 | 0.465471429 |
| Chord | 0.593805779 | 0.692689818 | 0.742408472 | 0.767321126 | 0.7925 | 0.394642857 |
| ChordP | 0.607758683 | 0.707027101 | 0.756869247 | 0.781819299 | 0.806785714 | 0.427428571 |
| cxds | 0.576279854 | 0.675031498 | 0.724828357 | 0.749788808 | 0.774785714 | 0.367342857 |
| doubletCells | 0.396983835 | 0.487445535 | 0.535271112 | 0.559821949 | 0.584685714 | 0.173985714 |
| DoubletDetection | 0.569370526 | 0.666356657 | 0.715610186 | 0.740426505 | 0.793114286 | 0.500857143 |
| DoubletFinder | 0.537830717 | 0.636466839 | 0.686221775 | 0.711172069 | 0.736171429 | 0.339428571 |
| Scrublet | 0.564203075 | 0.663581473 | 0.713449225 | 0.738419459 | 0.763414286 | 0.399771429 |
| Solo | 0.604236942 | 0.703224153 | 0.752987364 | 0.777930394 | 0.803142857 | 0.434714286 |

Furthermore, to thoroughly evaluate the performance of Chord and the four built-in methods under a wide range of doublet rates, we used random sampling to proportionally sample singlets and doublets in the dataset to build a doublet rate gradient (STAR methods), The performance of these methods generally showed an upward trend as the doublet rate increased, except that the DoubletFinder performance had dropped significantly at some doublet rates. In the gradient test of the DM-A dataset, bcds had the best performance while the effect of Chord was only slightly inferior to that of bcds. In the 15 datasets with doublet rates ranging from 2% to 30% generated from the HTO8 dataset, Chord ranked first in terms of the AUC on 10 datasets and the AUPRC on 9 data sets. (Figure S1C). Surprisingly, Chord outperformed better than other methods on many doublet rates.

### 2. The performance of doublet detection approaches on ground-truth datasets

In addition to the abovementioned computational doublet detection approaches based on the R environment, some cutting-edge doublet detection `software programs` based on the Python platform have been published in recent years. To integrate more doublet identification algorithms to improve the accuracy without losing the usability of Chord and the convenience of the R environment, Chord developed an expandable port for more doublet identification algorithms (Figure S1A). We can use this port to use the scoring results of other doublet detection software as input files, integrate these new methods with the four built-in methods and obtain a training model to further improve the accuracy of doublet identification. We used the Chord port to integrate three Python software programs (Scrublet(Wolock, Lopez and Klein, 2019), Solo(Bernstein, Fong, Lam, Roy, Hendrickson and Kelley, 2020) and DoubletDetection(DePasquale, Schnell, Van Camp, Valiente-Alandi, Blaxall, Grimes, Singh and Salomonis, 2019), Table S1) for an enhanced AdaBoost algorithm model, called Chord Plus version (ChordP).

To compare the doublet detection performances of Chord, ChordP and the other seven stand-alone `software programs`, we used the seven ground-truth scRNA-seq datasets mentioned above.Looking at the overall performance from the average of the seven ground-truth scRNA-seq datasets (Table S2), Chord and ChordP can achieve improved accuracy, and the more interesting part is the stability (Figure 2). Compared with Chord, the AUC of ChordP increased from 0.837 to 0.841 on the DM-A dataset and from 0.815 to 0.82 on the HTO8 dataset (Figure 2A), and the AUC value of ChordP on `datasets` other `than` DM-2.2 also had similar improvements. Remarkably, ensembling more methods can effectively improve the accuracy. On the other hand, the average AUC, pAUC800, pAUC900, pAUC950 and pAUC975 of ChordP across all the datasets were the highest among all methods (Table 1), and the average rank value in all datasets reached the highest (2.43, Figure 2B). These results showed that ChordP can indeed `obtain` more accurate results after ensembling 7 methods. In addition, the ranking variance of ChordP `was` 1.75, `which` was lower than `that of` Solo (3.05) and bcds (1.8), both of which had the same high accuracy rate (Figure 2C, Table S4). This finding shows that ChordP has better versatility for different datasets than the other methods. Through the uniform manifold approximation and projection (UMAP) method for `visualizing` the doublets, `singlets`, false negative (FN) results and false positive (FP) results (Figure S2), the distributions of the classification results between each method were quite different, which intuitively showed the complementarity between the different methods and the rationality of ensembling these methods. Among them, some methods, such as doubletDetection and DoubletFinder, had a concentrated spatial distribution of FP results in the HTO8 dataset. The removal of doublets based on these scoring results may lead to the accidental deletion of such FP cell-enriched clusters, which may affect the proportion statistics of cell types `and` may also directly lead to the deletion of rare cell `subpopulations`. In contrast, the FP results of Chord and ChordP were relatively evenly distributed and avoided becoming independent clusters and `affecting` subsequent analysis (Figure S2). Taken together, `the results` showed that ChordP`, which integrates` more algorithms, outperforms Chord and the other methods. We tested the time consumption of these different `software programs` under uniform hardware conditions and found that Chord did not significantly increase the time consumption. cxds was extremely time efficient, while Solo was time consuming in a CPU environment (Figure 2D).

**Figure 2.**
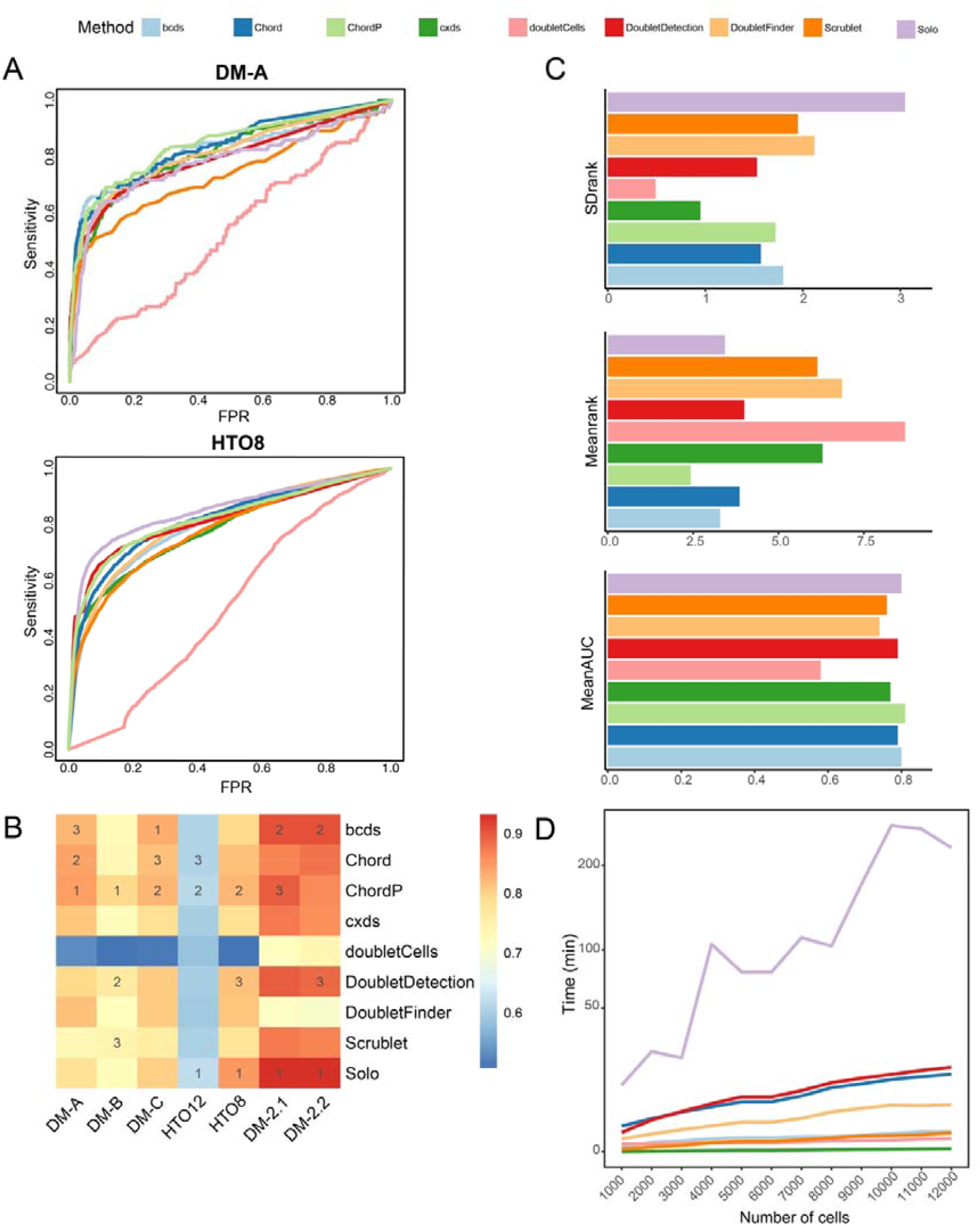
Comparison of the doublet detection approaches on the ground-truth datasets. (A) Nine methods evaluated on the DM-A and HTO8 `tests` and their `ROC curves`. (B) The `AUCs` of `the` nine methods on seven datasets and the heatmap of the `AUC results`. The number in the figure `indicates` the rank of the method in the dataset (only the top three methods are marked). (C) The standard deviation of the rank values for each method, the mean of the rank values of each method, and the mean AUC of each `method` across seven datasets. (D) By random sampling from the real dataset DM-2.1, simulated datasets with varying numbers of cell number (from 1,000 to 12,000 cells with `a`n interval of 1,000) were constructed, and the runtime of each method on the simulated datasets was recorded using the same computer server (E5-2678v3 CPU processor and `256 GB` memory).

### 3. The performance of doublet detection approaches in DEGs and pseudotime analysis

`To` evaluate the effect of different methods `on` downstream analysis, we used synthetic scRNA-seq datasets from recent benchmarking research (Xi and Li, 2020) to comapare Chord and other approaches in `terms` of differentially expressed gene (DEG) detection and pseudotime analysis. In the DEG analysis, one of the synthetic scRNA-seq `datasets was` ‘clean data’ with two cell types and 1126 between-cell-type `DEGs`, while the other dataset `was` the ‘contaminated data’ mixed with doublets at `a` 40% doublet rate (Figure 3D). We applied Chord and other approaches to remove the predicted doublets from the contaminated data by each method to obtain `a` ‘filtered dataset’ (Figure 3E). On the clean data, the contaminated data, and the filtered datasets by each doublet detection approach, DEGs were analysed using `the` Wilcoxon rank-sum test(Fay and Proschan, 2010) and model-based analysis of single-cell transcriptomics (MAST)(Finak, McDavid, Yajima, Deng, Gersuk, Shalek, Slichter, Miller, McElrath, Prlic et al., 2015). Three accuracy measures, namely, the true positive rate (TPR), true negative rate (TNR) and `precision`, were used to evaluate the `results` of DEGanalysis. In Figure 3A, `even` on the contaminated data, all the data processed by each doublet detection approach showed extremely high `precision` and TNRs on the two differential gene detection algorithms because these methods used more conservative methods to detect differential genes instead of detecting as many differential genes as possible so that the identified results `were` as correct as possible. `The` TPR (the percentage of the detected DEGs out of all the true DEGs) can better reflect the quality of the results. The low TPR for the contaminated data indicated that it was more difficult to identify DEGs in contaminated data, while the TPRs of the datasets processed by some doublet detection methods were higher than those for the clean data, which may be due to the deletion of transition state cells between cell types that were identified as doublets through these methods, resulting in obvious differences between cell types to detect more DEGs. In contrast, the differential gene analysis results of Chord and bcds were more similar to `those` on the clean data, indicating that the improved effect on differential expression analysis of `these` data using Chord and bcds was closer to the true situation.

**Figure 3.**
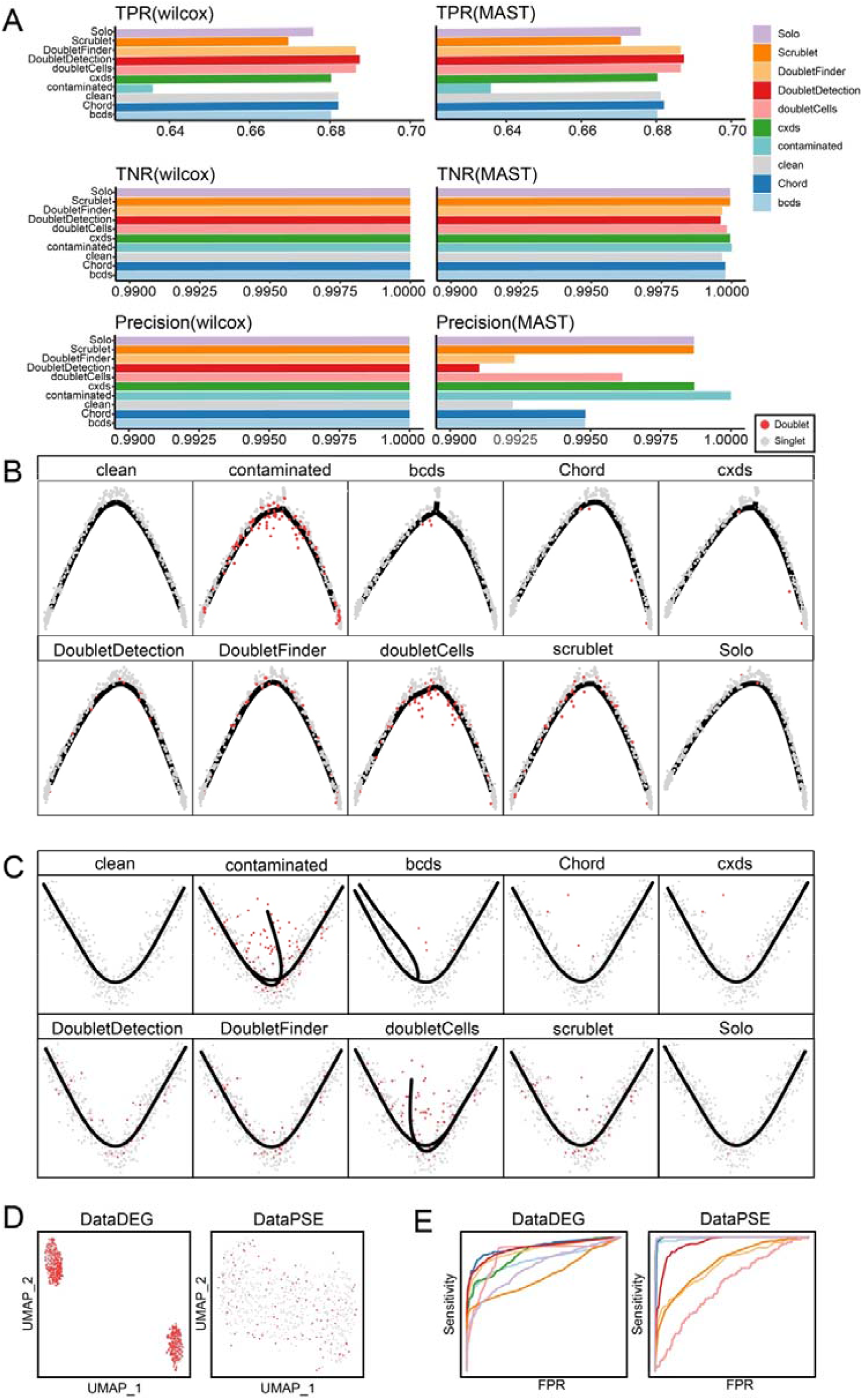
Evaluation of the doublet detection methods using the realistic synthetic datasets on DEG analysis and pseudotime analysis. (A) The dataset of labelled DEGs was processed by each doublet detection method, and the top 40% of cells based on the doublet score were excluded. Then, the DEGs were detected using MAST (Finak, McDavid, Yajima, Deng, Gersuk, Shalek, Slichter, Miller, McElrath, Prlic et al., 2015) and Wilcoxon rank-sum tests (Fay and Proschan, 2010). Taking the doublet as positive, three accuracy measures (i.e., the TPR, TNR and precision) were calculated. (`B, C`) After processing the dataset for the pseudotime analysis using each doublet detection method, the top 20% of cells according to the doublet score were excluded. Monocle 2 `(B) Slingshot (C)` were used to construct the trajectories of these results. (D) The UMAP was embedded for the two realistic synthetic datasets (DataDEG and DataPSE), in which the doublets are shown in red and the singlets are shown in grey. DataDEG is a simulation dataset containing two synthetic cell types, including 1,667 cells, 40% of which are correctly labelled doublets. DataPSE consists of 600 cells, 20% of which are synthetic labelled doublets containing a bifurcating trajectory. (E) The AUC of each method on DataDEG and DataPSE and their ROC curves.

In the pseudotime analysis, the ‘clean data’ is a synthetic scRNA-seq dataset including a bifurcating trajectory, while the ‘contaminated data’ is composed of ‘clean data’ plus 20% doublets (Figure 3D). The ‘filtered dataset’ was generated by using Chord and other approaches to remove the predicted doublets from the contaminated data (Figure 3E). Then, two pseudotime analysis methods (Slingshot (Street, Risso, Fletcher, Das, Ngai, Yosef, Purdom and Dudoit, 2018) and Monocle2 (Qiu, Mao, Tang, Wang, Chawla, Pliner and Trapnell, 2017)) were implemented on the clean data, the contaminated data, and the filtered datasets. In Figure 3B, using Monocle 2 `to infer the` cell trajectory, the contaminated data and the filtered datasets of bcds and cxds generated additional bifurcation `trajectories` due to the influence of doublets. The unrecognized real doublets by doubletCells had a clear tendency to deviate from the trajectory distribution. In the results of Slingshot (Figure 3C), the trajectories of the contaminated data and the filtered datasets of bcds and doubletCells obviously had one more fork. In contrast, Chord and Scrublet had similar cell `trajectories` to the clean data in the two pseudotime analysis methods, and there were fewer remaining doublets and no new branches were generated. Thus, we can see that Chord was equivalent to or even outperformed other methods in DEG detection and pseudotime analysis on the synthetic scRNA-seq datasets.

### 4. Applying Chord to real-world scRNA-Seq data

To take a closer look at the application of Chord in real-world data and whether the effect of downstream analysis has been improved, we tested the Chord method on a real-world scRNA-seq data dataset without doublets labelling information that containing 52,698 cells from lung cancer tumour tissues of 5 patients (Lambrechts, Wauters, Boeckx, Aibar, Nittner, Burton, Bassez, Decaluwé, Pircher, Van den Eynde et al., 2018). Based on the expected doublet rate (0.9% per 1,000 cells), we estimated the proportion of doublets in 5 patient samples (2.57%, 1.43%, 11.88%, 15.01%, and 16.65%), and the number and proportion of doublets for different cell types. After removing doublets predicted by Chord, we can see that T cells contained the largest number of doublets due to their large cell number. The identified doublets of B cells accounted for only 7.6%, while the number of doublets of epithelial cells was the smallest, but the proportion was the highest (39.25%) (Figure 4A). On the UMAP plots (Figure 4B), doublets were unevenly distributed, clustered at the edge of some clusters, and even formed independent clusters. Obviously, doublet removal by Chord can have a greater impact on the proportion of cells to avoid these imbalanced distributions and numerous doublets from becoming noise contamination for the quantitative statistics of the proportion of cell types.

**Figure 4.**
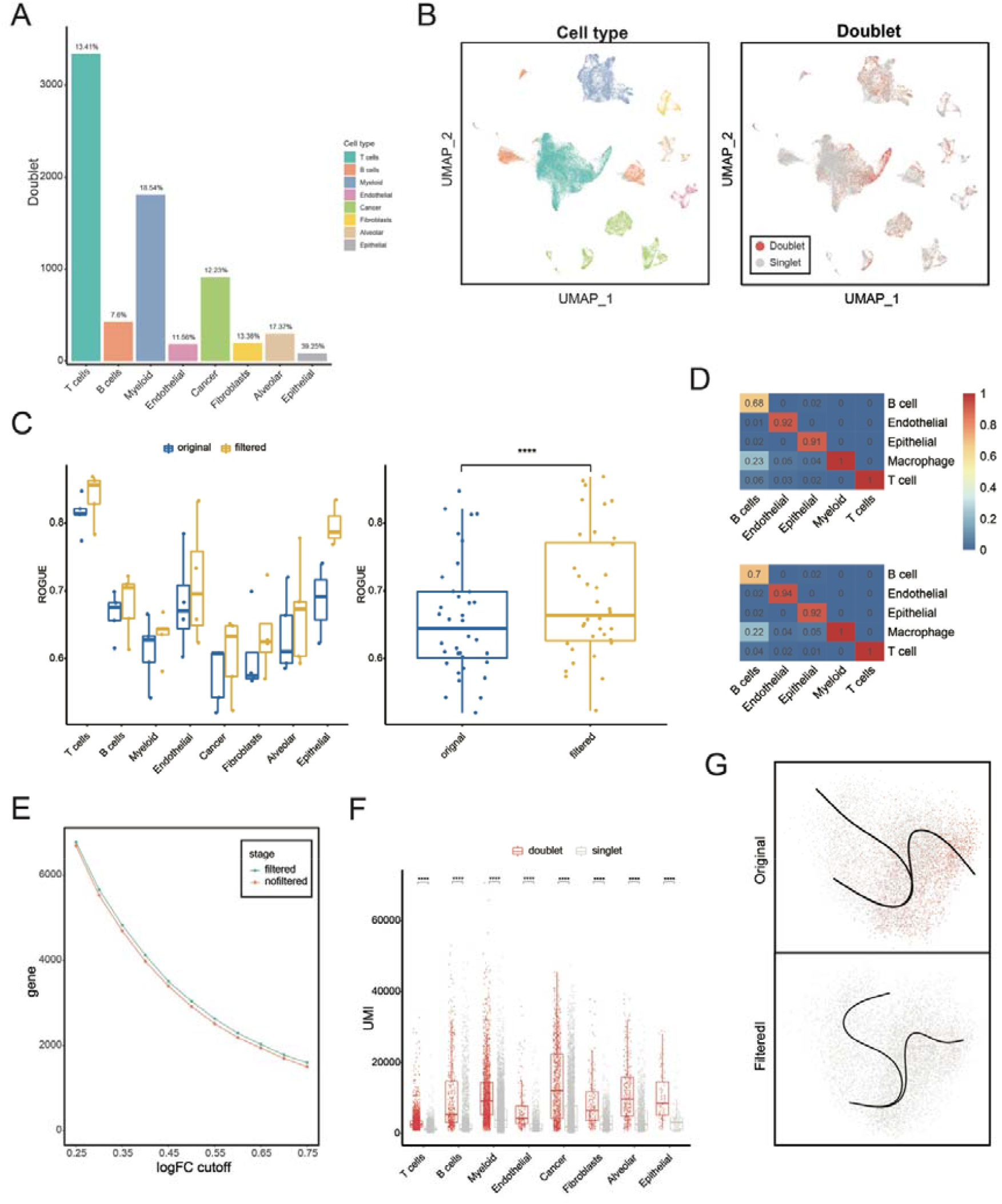
Doublet removal by Chord improves the analysis performance on real-world scRNA-Seq data. (A) `Doublet detection was performed` on the published lung cancer data set (Lambrechts, Wauters, Boeckx, Aibar, Nittner, Burton, Bassez, Decaluwé, Pircher, Van den Eynde et al., 2018) using Chord, and recorded the number and the proportion of `doublets` for each cell type `were recorded`. (B) UMAP on the 52,698 cells of this dataset. The cells were coloured by cell type (left). Each cell was coloured by `the` predicted result of doublet detection (right). (C) The ROGUE value (Liu, Li, Li, Wang, Ren and Zhang, 2020) of each cell type for each sample. `A paired` t-test was used to test the difference `in` each cell type in each sample between the two groups before and after `doublet` filtration (paired t-test, * p<0.05, ** p<0.01, *** p<0.001, **** p<0.0001). (D) Automated cell type annotation of this data set was `performed` using SciBet (Li, Liu, Kang, Liu, Liu, Chen, Ren and Zhang, 2020). The training model provided by SciBet, which was trained from 42 human single-cell datasets containing 30 major human immune cell types, was used to automatically annotate the datasets before and after doublet `removal`. `The` matrix heatmaps are plotted. (E) A line chart `showing` the changes `in` the total number of differentially expressed genes before and after doublet `removal`. The DEGs were calculated by the Wilcoxon rank-sum test (Seurat). The threshold value of logFC was measured by a gradient from 0.25 to 0.75 at 0.05 intervals. (F) The RNA UMI numbers `were significantly different` between the doublets and singlets predicted by Chord in each cell type (unpaired t-test, * p<0.05, ** p<0.01, *** p<0.001, **** p<0.0001). (G) The myeloid cells in original and filtered data for the pseudotime analysis were processed using Slingshot (Street, Risso, Fletcher, Das, Ngai, Yosef, Purdom and Dudoit, 2018). The trajectory of the original data and the filtered data are shown respectively.

To evaluate the performances of Chord on real data to improve the effectiveness of downstream analysis. We first calculated the ratio of global unshifted entropy (ROGUE) value (Liu, Li, Li, Wang, Ren and Zhang, 2020) representing the purity of a cell type or cluster on the original data and filtered data after the doublets `were removed and` processed by Chord. As a result, the ROGUE value of the filtered data was significantly improved, although for each cell type there was not enough power to detect a significantly improvement, but the increasing trend of ROGUE value after filtering doublets was obvious (Figure 4C). Second, we used the cell type annotation software SciBet(Li, Liu, Kang, Liu, Liu, Chen, Ren and Zhang, 2020) to build a model based on real data to annotate the cell types of the immune system cells in the original data and filtered the data to examine the changes in the differentiation of cell types before and after applying the doublet filter. The accuracy of the annotation results of the B cells (0.7), `endothelial cells` (0.94), `and epithelial cells` (0.92) on the filtered dataset, `were` all greater than those of the B cells (0.68), `endothelial cells` (0.92), `and epithelial cells` (0.91) in the unfiltered dataset (Figure 4D). It is worth noting that cell purity will also affect the calculation of DEGs between different subgroups. Doublet `simulation` from multiple cell types will make it more difficult to find DEGs (Figure 4E). Since `a` doublet is caused by multiple cells with the same barcode, doublet cells generally contain `a` higher unique molecular identifier (UMI). In the doublet detected by Chord, the UMI of all cell types `was` significantly higher than that of other cells (Figure 4F). A pseudotime analysis `of` the myeloid cells was conducted. The doublets were unevenly distributed in the dimensionality reduction results and `aggregated on` the right side. Since the trajectory analysis was based on the fitted trajectory, the doublets far away from trajectory would lead to a deviation. In the pseudotime analysis after filtering the doublets, the direction of `the` trajectory changed (Figure 4G). We infer that the deviation was corrected by removing the `doublet` data.

Therefore, we believe that Chord’s doublet processing of real data can improve the purity of cell populations, allowing researchers to obtain more accurate cell type identification results, accurately identify DEGs between cell types and obtain better pseudotime analysis result.

## Discussion

At present, there are a variety of doublet detection methods, and these methods perform very `differently`. On the one hand, different methods are specific to different data sets (Figure 2B). On the other hand, for the same data set, the distribution of TP, FP and FN results identified by different methods is significantly different (Figure S2A, Figure S2B). For users, it is difficult to evaluate and choose the most suitable method. To solve the problem, Chord, which integrates the results of different methods through the AdaBoost algorithm, takes the essence of the methods and minimizes the disadvantages. According to the evaluation, Chord, with high average ranking and stability, is widely applicable to various data sets. Chord is also scalable and can be integrated by simply importing the cell’s evaluation scores of any method. `An increasing number of` better methods can be integrated to further improve accuracy (Figure 2B, Figure 2C). It will be compatible with any new approach in the future, and it can accept any update to the existing approaches (Figure S1A).

Doublets contained in scRNA data affect not only the quantity of cell types but also the accuracy of downstream analysis. The data filtered by Chord `were` more `accurately` identified in cell `type` annotation (Figure 4D) and `tended` to find more possible DEGs (Figure 4E). `This` phenomenon would help researchers `obtain` more accurate results and `conclusions in subsequent analyses`. Unfortunately, Chord is not optimal in terms of time efficiency, and perhaps we will improve this with further refactoring of the integrated methods in the future. `In addition`, doublet detection methods and other `pre-processing methods,` such as removal of ambient mRNA contamination in `droplets,` may `interact` with each other(Yang, Corbett, Koga, Wang, Johnson, Yajima and Campbell, 2020). How `the` pre-processor `affects` the different types of methods and how to `obtain` better results with other `pre-processing methods` is worthy of detailed evaluation in the future.

We have provided an improved idea that optimized a general step of most `doublet` detection methods, constructing `a` simulated training set. Previous methods add simulated doublets to real data and regard it as `a` training set. Although the simulated doublets were labelled correctly, a large number of real doublets `that` were undiscovered were labelled incorrectly as singlets. `This` resulted in `an` inaccurate training set and affected models. `Correspondingly`, Chord `added` a step, ‘overkill’, which first used different methods to evaluate the data, filtered out cells identified by any method, `and` then simulated doublets by the remaining cells. In this way, the doublets mislabelled as singlets in the training set were greatly reduced, thus `improving` the labelling accuracy of the training set and the AUC value of the final result. For methods `such` as bcds, DoubletFinder, and DoubletDetection, which use random sampling from real data to simulate doubles and construct training sets, this idea works. Overall, we proposed a computational approach for doublet detection called Chord that uses an ensemble algorithm model to classify cells. This is the first `study` of its kind using `an` ensemble algorithm on doublet detection. This work `helps` researchers save time and energy `in` selecting and testing individual software `for` doublet detection, solve this problem in a more convenient way, and `obtain` better results. We expect Chord to become an essential part of the future scRNA analysis process, and we have shared `the` source code of Chord on GitHub.

## Acknowledgment

We thank Dr. Yong Bai from BGI-Shenzhen for the algorithm suggestions. This research was supported by the Guangdong Enterprise Key Laboratory of Human Disease Genomics (2020B1212070028), Shenzhen Key Laboratory of Single-Cell Omics (ZDSYS20190902093613831).

## Author Contributions

H.L.Z and K.X.X designed the research, performed the data analyses, and wrote the codes and the manuscript K.K, J.H.Y, H.M.Y suggested optimization to the article. G.B.L provided critical advice and oversight of the research and revised to the manuscript. All authors have read and approved the final manuscript.

## STAR Methods

### KEY RESOURCES TABLE

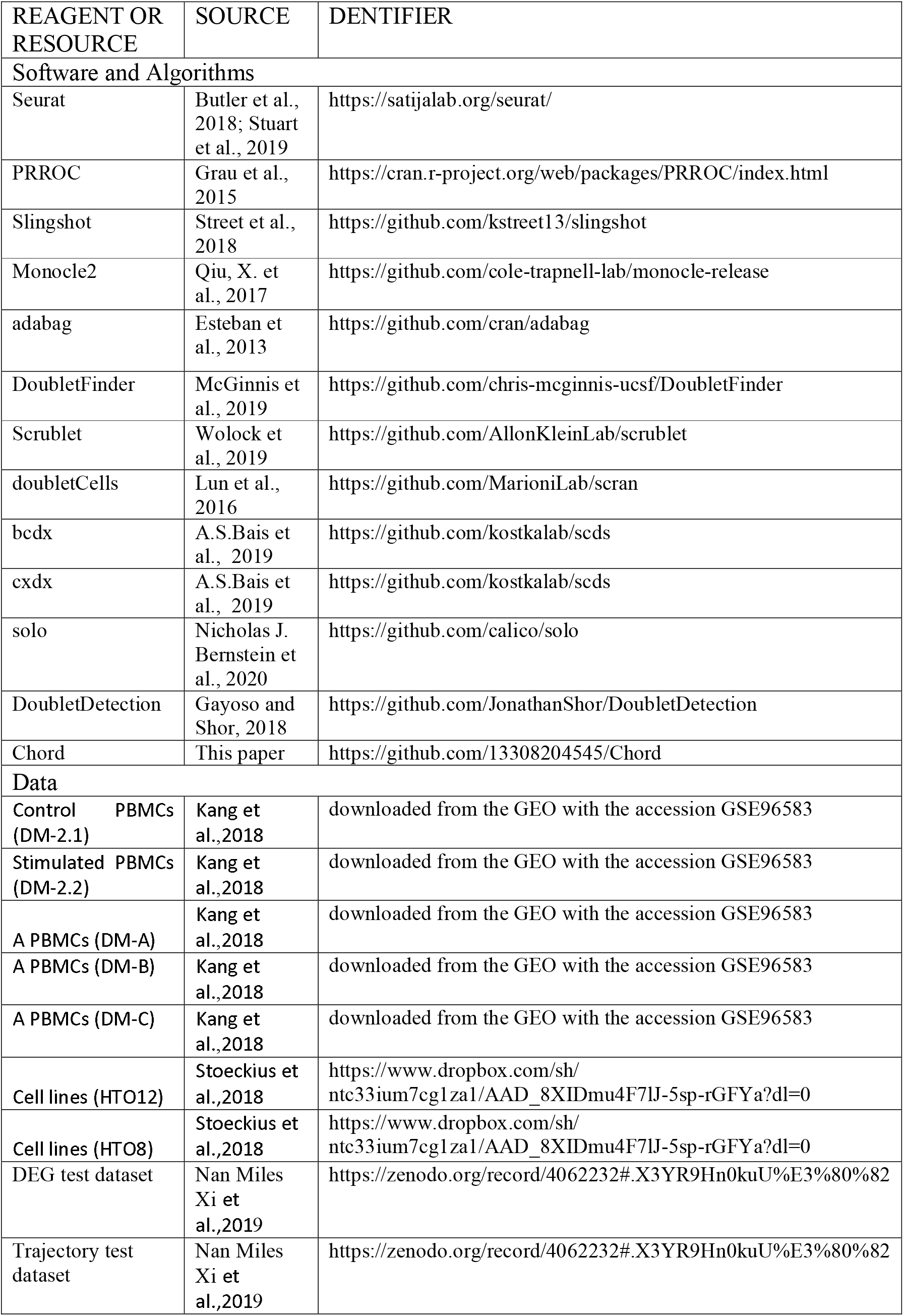

## CHORD OVERVIEW

### Data input

Chord’s input format is `a` comma-separated expression matrix `that` is a background-filtered, UMI-based `count` matrix of a single sample. For the `count` expression matrix, Chord will pre-process it according to the Seurat analysis pipeline. Chord can also directly accept object files generated by the Seurat analysis pipeline. In addition, it is suggested `that` users estimate a doublet rate (Figure S1) based on the loading conditions so that Chord can simulate a simulated training set `that` is closer to the real dataset. It should be mentioned that the doubletrate, which is the estimated doublet rate parameter, has a certain degree of robustness, and a certain degree of deviation will not `greatly` affect the result of Chord.

### Data pretreatment

`To` train the results of bcds, cxds, DoubletFinder, and DoubletCell through ensemble learning, Chord created SingleCellExperiment object data conforming to the input format of bcds and cxds through the R package SingleCellExperiment(Amezquita, Lun, Becht, Carey, Carpp, Geistlinger, Marini, Rue-Albrecht, Risso, Soneson et al., 2020).

### Preliminary deletion of doublets

We tend to reduce the number of doublets in the sample as much as possible before generating simulated doublets to avoid the datasets `that are` prepared for model training containing real `doublets` because real doublets were not labelled and are labelled as singlets in the training set. We `applied` four doublet detection methods, the `dataset` was evaluated, and the `doublets were` roughly filtered out.

We used the scds() function, cxds, and bcds to evaluate the dataset with parameters ntop = 500, binThresh = 0, retRes = T, and extracted the doublet scores evaluated by two methods for each cell; through the DBF() function in the called R package DoubletFinder, used the no ground-truth process given by the author for evaluation, the parameter selection was PCs = 1:10, pN = 0.25, and automatically extracted the pk value corresponding to the highest bimodal coefficient, and extracted the doublet score; doubletCells() was used to evaluate doublet cells with parameters k = 50, d = 50 and extract the scores.

Chord introduced an adjustable parameter” overkillrate. According to this ratio, we filtered the doubletrate*overkillrate percentage of cells that were most likely to be doublets according to the evaluation results of each method and `obtained` the ‘pre-filtered data’.

### Generating the simulation data set

In the case of avoiding the generation of `doublets` synthesized by the same cell type, Chord randomly sampled `pairs` of cells in the pre-filtered data, generated simulated doublets from the raw UMI count by mixing the gene expression profiles of the selected cell pair(Wolock, Lopez and Klein, 2019) and then added simulated doublets to the pre-filtered data:

1. Perform a Seurat standardization process on the data; call the functions NormalizeData(), FindVariableFeatures(), ScaleData() with default parameters.

2. Cluster cells, perform dimensionality reduction operations on the data through RunPCA(), taking PC1 to PC30 as input, and perform `k-means` clustering (k=20). After clustering, divide the cells into 20 clusters.

3. Randomly sample pair of cells at a ratio of doubletrate/(1-doubletrate) for each cell type, and weight the cells with introduced biological random number from a N(1, 0.1) distribution.

4. Average the weighted gene expression profiles of the cell pair as simulated `doublets`.

5. Add the simulated doublets to the pre-filtered data; take this new dataset as the training set.

### Model training

For the training set where all cells were labelled, Chord used the same parameter settings to evaluate the doublet scores through the bcds, cxds, DoubletFinder, and DoubletCell methods. `Using` the R package ‘adabag’(Alfaro, Gamez and García, 2013), through `the` AdaBoost algorithm, DBboostTrain() `was used` to implement model training `for training` based on the scoring results of these four methods. The number of iterations `was` set to 40, the simulated doublets were set as `true positives (TPs),` and the singlets were set as `true negatives (TNs`) for model training.

### Scoring the original data

We called DBboostPre(), used the model trained on the training set to predict the doublet scores of the original dataset, and output the doublet score of each cell based on the result of our ensemble model.

### Expandable interface

In response to possible version updates of integrated doublet tools and the release of new doublet tools, Chord includes an extendible interface that can be customized based on the doublet evaluation results of any tool. The chord() function can extract the expression matrix after generating the simulated doublets. `The user` needs to export pre-filtered data through the chord() function, `evaluate` the original data and pre-filtered data using the intended methods, `and` then `import` the scores into Chord. Based on the evaluation scores, Chord uses the AdaBoost algorithm to ensemble the extra methods together with bcds, cxds, DoubletFinder, and DoubletCell `and` then `uses` the ensemble model to score the original dataset.

### Solo settings

Solo was run in the Python 3.7.9 environment, and the parameters were set according to the reference file solo_params_example.json downloaded from https://github.com/calico/solo. The doublet scores of each cell was read through the softmax_scores.npy file.

### Doublet detection settings

`Double detection was run in the Python 3`.7.9 environment using the operating parameter settings from https://nbviewer.jupyter.org/github/JonathanShor/DoubletDetection/blob/master/tests/notebo oks/PBMC_8k_vignette.ipynb. Obtain scoring results through BoostClassifier.fit.voting_average_.

### Scrublet settings

Scrublet was run in the Python 3.7.9 environment, following the `instructions at` bhttps://github.com/AllonKleinLab/scrublet/blob/master/examples/scrublet_basics.ipynb. The doublet scores of each cell was read through Scrublet.scrub_doublets.

### Doublet rate gradient

The doublet gradient dataset `is` composed `of` random sampling from real datasets. The dataset HTO8 is randomly sampled from the dataset DM-A with an interval of 0.02, and the ratio of the doublet rate is 0.02 to 0.30. The dataset DM-A is randomly sampled from the dataset DM-A with an interval of 0.01, and the ratio of the doublet rates is 0.01 to 0.10.

### Method rank variance comparison

In view of the unstable performance of the method on different datasets, we calculated the AUC rank variance coefficient to characterize the stability of the method on different datasets. For the methods participating in the evaluation, rank according to their AUC, and calculate the variance of the AUC ranking of each method under different datasets as the AUC rank variance coefficient (SDrank). The method with the highest SDrank is unstable on different datasets, where the SDrank represents the generalizability of the method.

## QUANTIFICATION AND STATISTICAL ANALYSIS

### AUROC, AUCPR calculation

Each `method` was evaluated using `the` AUC with ground-truth labels of original datasets or labels of simulated datasets. `In addition, the` AUPRC `and` partial area under the ROC curve (pAUC) were calculated. We calculated the AUC and AUPRC by means of the PRROC R package(Grau, Grosse and Keilwagen, 2015), and plotted the ROC curve of single tools by setting the option ‘ROC curve’ to ‘TRUE’. For the pAUC at 0.9, a 0.95 specificity `was` calculated by using the ‘pROC’(Robin, Turck, Hainard, Tiberti, Lisacek, Sanchez and Müller, 2011) R package.

### Evaluating DEGs with simulated datasets

We used the published simulated single-cell sequencing dataset (Xi and Li, 2020) to test the changes in the number of `DEGs` correctly detected before and after doublet removal by different methods. The simulated single-cell dataset was generated by scDesign (Li and Li, 2019) and contained 1,667 cells and 18,760 genes. It was divided into two cell types, 500 cells and 667 doublets. Among them, high-expression and low-expression DEGs, which were known at the time of data generation, accounted for 6% (3% `upregulated` genes and 3% `downregulated` genes). The dataset without doublets was used as the clean dataset, and the data with doublets were added as the contaminated dataset. After the contaminated dataset was evaluated by a doublet detection method, the dataset of 40% of the cells with the highest score `was` filtered according to the result. We performed the process described above for each method. After that, we used Seurat’s FindAllmarkers() function with the methods ‘wilcox’(Fay and Proschan, 2010) and ‘MAST’(Finak, McDavid, Yajima, Deng, Gersuk, Shalek, Slichter, Miller, McElrath, Prlic et al., 2015) to perform `DEG` calculations on the dataset. `Finally,` we calculate the precision, TPR, `and` TNR for the data for all of the datasets.

### Pseudotime analysis of simulated data

We used the published dataset of simulated single-cell sequencing(Xi and Li, 2020) to test the effects of different methods on pseudotime analysis. The simulated single-cell dataset, generated by Splatter, consists of 600 cells and 1,000 genes. There are two cell tracks containing 250 simulated cells and 100 simulated doublets. Taking the dataset without the doublets as a clean dataset, and the data containing the doublets as a contaminated dataset. After evaluating the contaminated data set through a doublet detection method, the highest scoring cells for the number of double cells according to the results were removed. After Then, the R package Monocle and Slingshot software were used for pseudotime analysis of the dataset. We performed the process described above for each method. `In addition`, we calculated `the trajectories of the` clean and contaminated datasets.

### Time cost

By random sampling from the real dataset DM-2.1, we constructed test datasets of 1,000 to 12,000 cells with `a` gradient of 1,000 cells and tested various methods with the same processor. Then, the runtime of each method on the simulated datasets `on` the same computer server (E5-2678v3 CPU processor and `256 GB` memory) `was recorded`.

### Base analysis process of lung cancer data

First, a Seurat standardization process was performed on the data, the functions NormalizeData(), FindVariableFeatures(), ScaleData() were called with default parameters, and then dimensionality reduction was performed on the data through RunPCA(). Then, we `constructed a shared nearest neighbour (SNN) graph` for a given dataset with PC1 to PC30 by FindNeighbors(), and we `identified` clusters of cells by an SNN modularity optimization`-` based clustering algorithm with resolution=0.5. We computed the UMAP embeddings, which included annotated experimental doublets and `cell types` annotated by markers provided by the original data(Lambrechts, Wauters, Boeckx, Aibar, Nittner, Burton, Bassez, Decaluwé, Pircher, Van den Eynde et al., 2018)

### Cluster purity

We calculated the ROGUE value for each `cell type` in each sample. Comparisons between two original `datasets` and filtered `datasets` were `performed` using paired two-tailed t-tests. Filtered or not was the grouping condition. The environment parameters were platform = ‘UMI’ and span = 0.6.

### Cell type identification using SciBet

We used SciBet(Li, Liu, Kang, Liu, Liu, Chen, Ren and Zhang, 2020) to perform cell type analysis on epithelial, endothelial, myeloid, T, and B cells before and after processing. The identification reference is the reference set of human main single`-`cell sequencing cell types provided by SciBet (http://scibet.cancer-pku.cn/major_human_cell_types.csv).

### Calculated number of DEGs

The number of DEGs were calculated by the Wilcoxon rank-sum test (Seurat) with the following parameters: min.pct = 0.1, test.use = ‘wilcox’, and the gradient logfc.threshold from 0.25 to 0.75 at an interval of 0.05. Then, we counted the number of DEGs at different logfc.threshold values.

## Supplemental Information

### Supplemental figure

**Figure S1.**
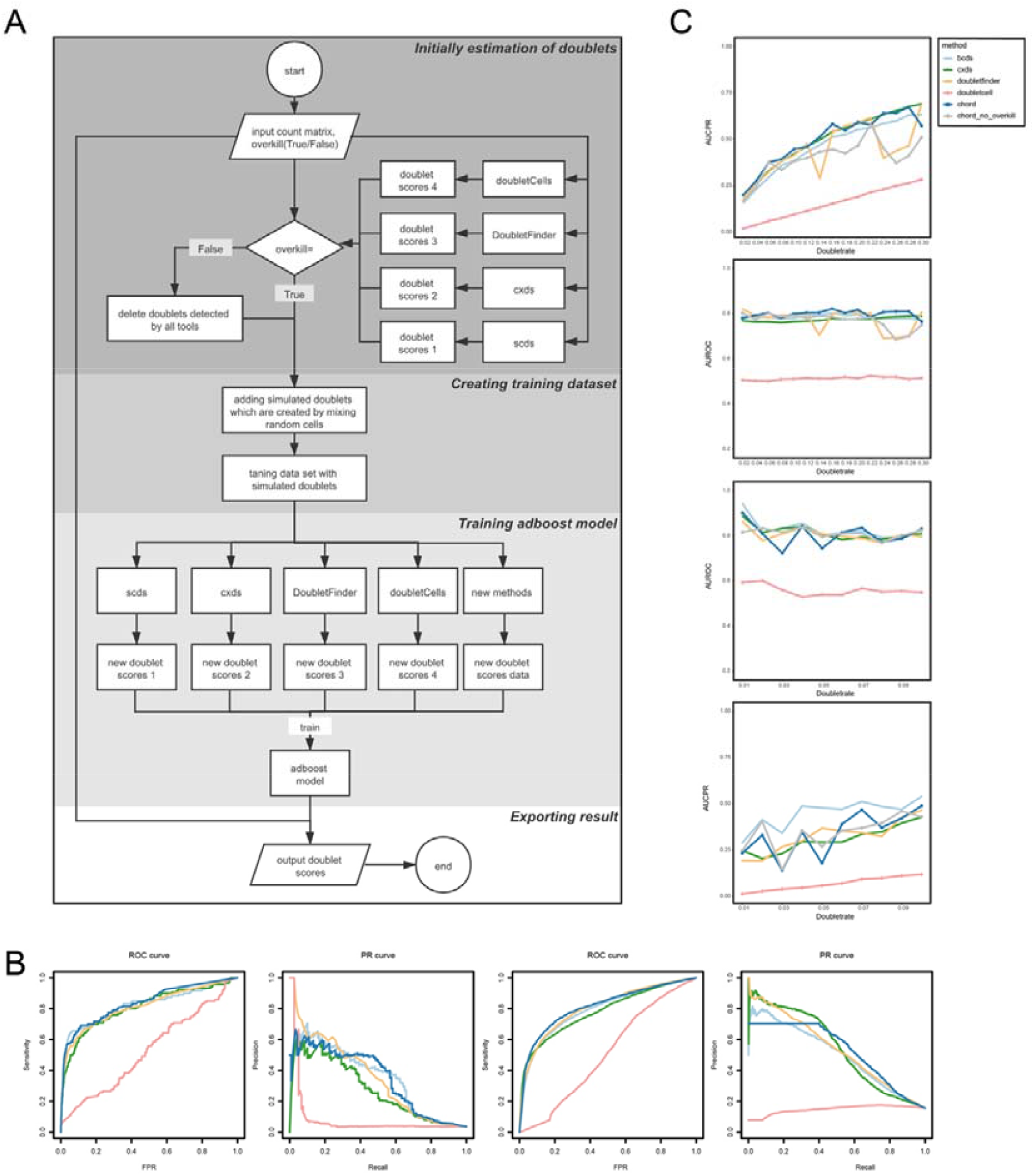
The flow chart of Chord. (A) The workflow of the Chord software is divided into four parts. The flow chart shows the data flow of the software. (B) The receiver operating characteristic curves (ROC) and PR curve of Chord and the other four methods were draw for the test DM-A (the first row) and HTO8 (the second row). (C) We generated the HTO8 sub-datasets with doublet rate from 0.02 to 0.30 (the first two from the top) by randomly sampling the dataset, and the sub-datasets with doublet rate from 0.01 to 0.10 of the data set DM-A (the third and fourth from the top). The values of AUC and PAUC on these datasets are calculated respectively for Chord and the other four methods.

**Figure S2.**
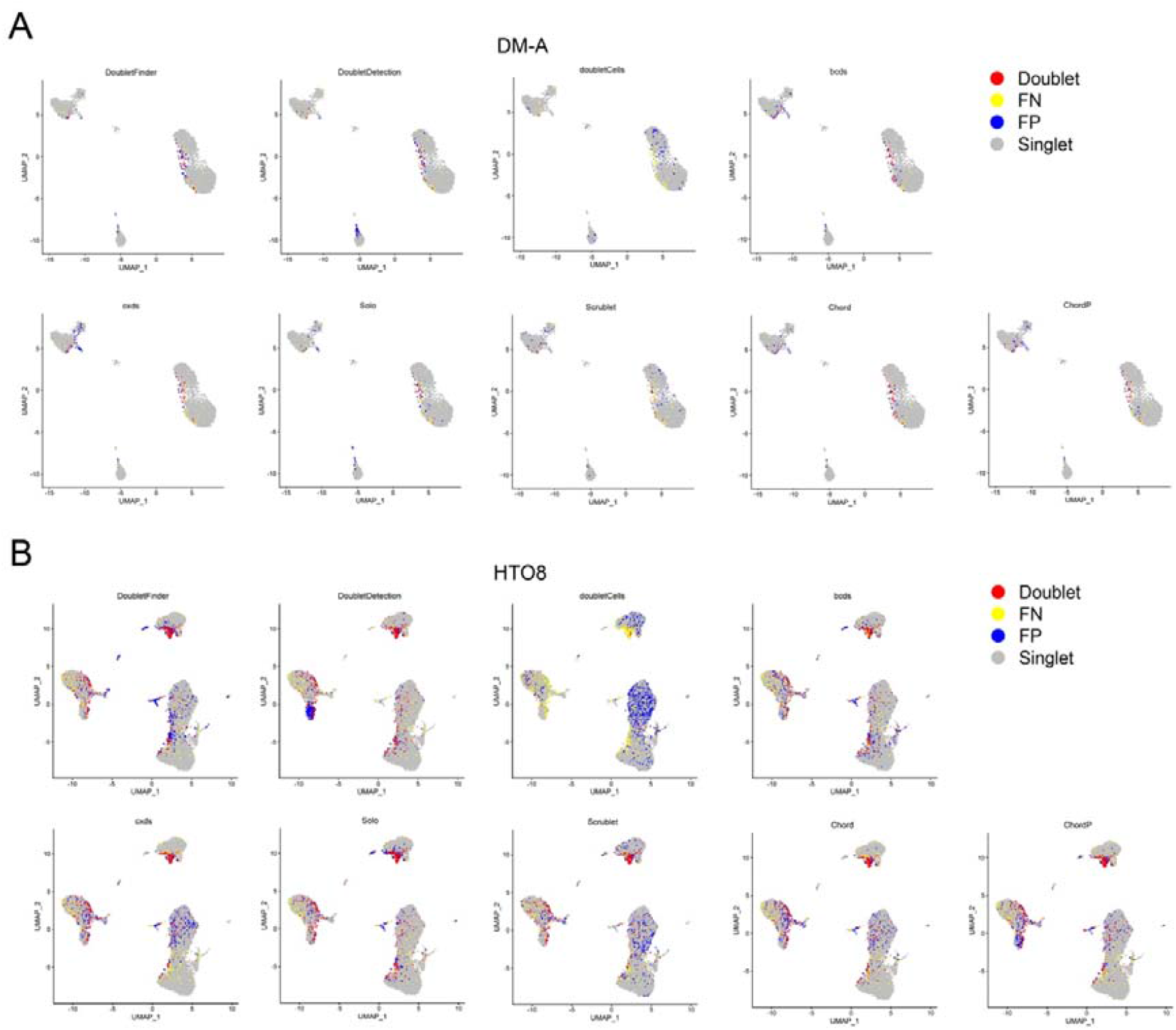
The UMAP visualization of dataset DM-A and dataset HTO8 on doublet, singlet, FP and FN. For the real dataset DM-A (Fig.S2 A) and dataset HTO8 (Fig.S2 B), according to the scoring results of each method, we deleted the top scoring cells according to the real doublet rate, and verified with the real label to visually show the doublet, singlet, false positive (FP) and false negative (FN) results.

## Supplemental table

table S1 Overview of the computational methods for doublet detection and their capabilities

table S2 Overview of the real scRNA-seq datasets with experimentally annotated doublets used in the study

table S3 Values of PAUC&PR for all methods in Table1 in each dataset

table S4 Rank values related information in Fig 2C

table S5 Time-consumption based on cell gradient

table S6 AUC corresponding to DM-A dataset in Fig S1B

table S7 AUC corresponding to HTO8 dataset in Fig S1B

table S8 PR corresponding to DM-A dataset in Fig S1B

table S9 PR corresponding to HTO8 dataset in Fig S1B

