## Supplementary figures and images for "Chord: Identifying Doublets in Single-Cell RNA Sequencing Data by an Ensemble Machine Learning Algorithm"

### Supplemental Figure 1

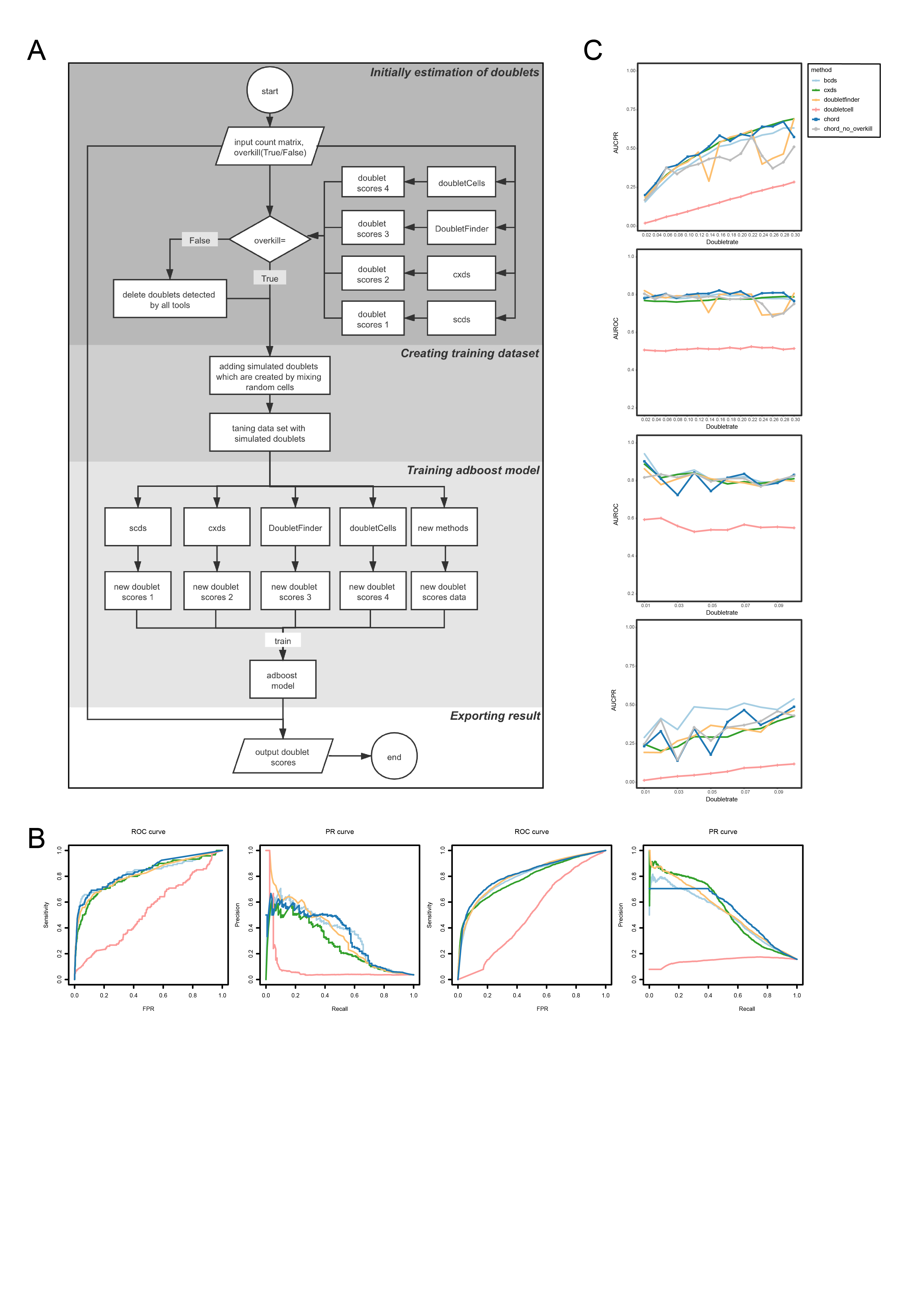

### Supplemental Figure 2

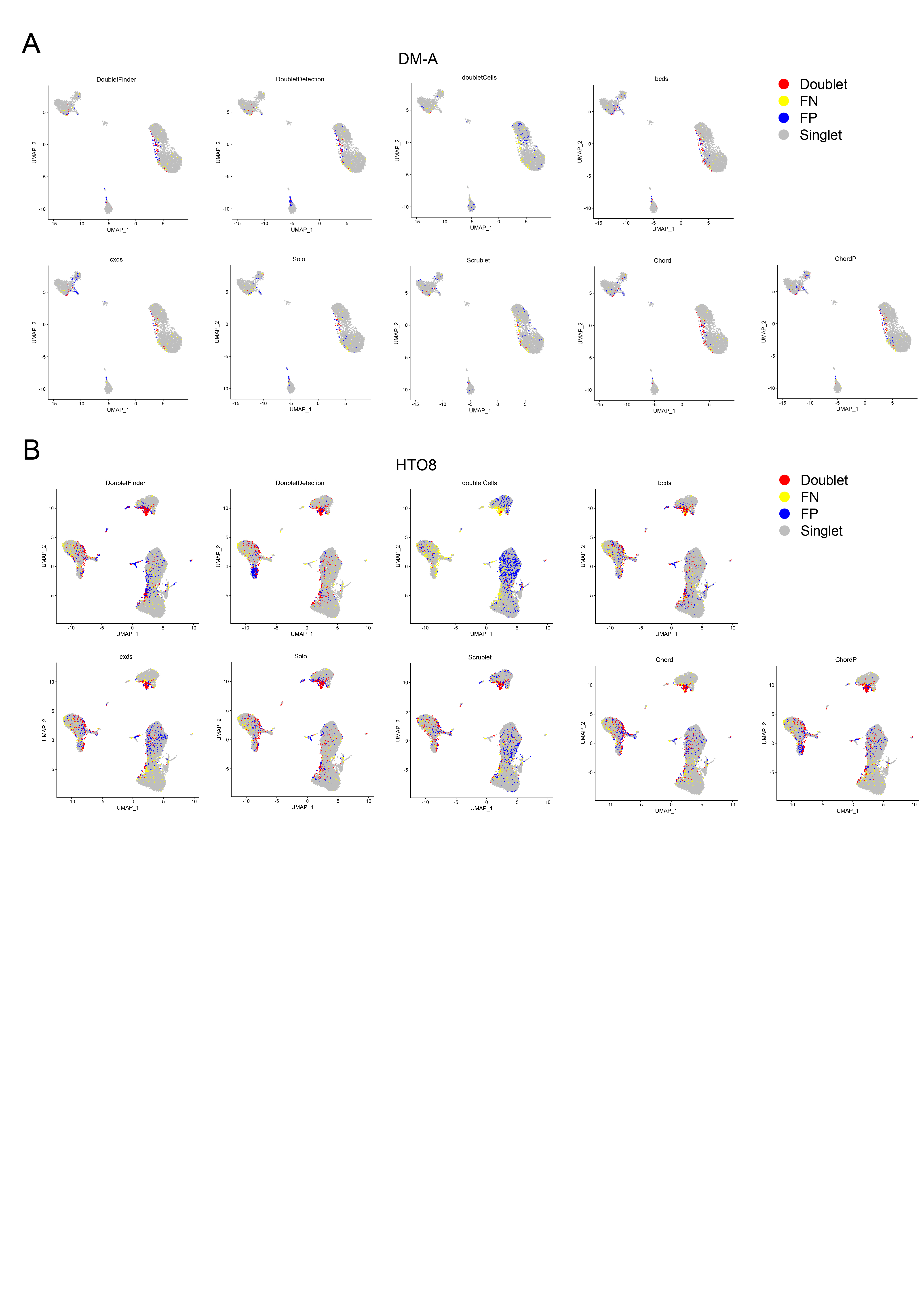
